# Long Noncoding RNA- Maternally Expressed Gene 3 Contributes to Hypoxic Pulmonary Hypertension

**DOI:** 10.1101/243063

**Authors:** Yan Xing, Xiaodong Zheng, Yao Fu, Jing Qi, Minghui Li, Haisheng Peng, Shuang Wang, Shuzhen Li, Daling Zhu

## Abstract

The expression and function of long noncoding RNAs (lncRNAs) in the development of hypoxic pulmonary hypertension, especially in the proliferation of pulmonary artery smooth muscle cells (PASMCs) are largely unknown. Here, we characterized the expression of lncRNA-maternally expressed gene 3 (lncRNA-MEG3) was significantly increased and primarily located in the cytoplasm of PASMCs by hypoxia. LncRNA-MEG3 knockdown by lung-specific delivery of small interfering RNAs (siRNAs) significantly prevented the development of hypoxic pulmonary hypertension *in vivo*. Silencing of lncRNA-MEG3 by siRNAs and gapmers attenuated PASMC responses to hypoxia *in vitro*. Mechanically, we found that lncRNA-MEG3 acts as a molecular sponge of microRNA-328 (miR-328); upon hypoxia, lncRNA-MEG3 interacts and sequesters miR-328, leading to the upregulation of insulin-like growth factor 1 receptor (IGF1R). Additionally, higher expression of lncRNA-MEG3 and IGF1R, and lower expression of miR-328 were observed in PASMCs of iPAH patients. These data provide insight into the contribution of lncRNA-MEG3 in hypoxia pulmonary hypertension. Upregulation of lncRNA-MEG3 sequesters cytoplasmic miR-328, eventually leading to the expression of IGF1R, revealing a regulatory mechanism by lncRNAs in hypoxia-induced PASMC proliferation.

## INTRODUCTION

Pulmonary hypertension (PH) is a severe disease resulting in right heart failure and even death (1). Hypoxia is a common cause of PH in multiple diseases, such as chronic obstructive pulmonary disease, sleep apnea and high-altitude pulmonary vascular disease, which primarily includes hypoxic pulmonary hypertension (HPH) (2,3). HPH is characterized by the elevation of pulmonary arterial pressure and hypoxia-induced pulmonary arterial remodeling (HPVR). Although many factors have been implicated in the susceptibility to HPVR, it is well-accepted that HPVR is primarily caused by excessive proliferation of pulmonary artery smooth muscle cells (PASMCs) (4). However, the exact mechanism regulating the pathological process of HPVR is largely unknown (3).

The roles of noncoding RNAs in HPH have been evaluated in many studies. MicroRNAs (miRNAs) are short noncoding RNAs associated with PH onset, progression, or treatment responsiveness (5). Our previous study showed that microRNA-328 (miR-328) by targeting the expression of insulin-like growth factor 1 receptor (IGF1R) and L-type calcium channel-alpha 1C is important in the development of HPH (6). However, the specific regulation of miR-328 expression in PASMCs is unknown. Long noncoding RNAs (lncRNAs) are a class of noncoding RNAs longer than 200 nucleotides and function in diverse biological progresses (7–11). Importantly, lncRNAs interact with microRNAs to modulate cell differentiation, proliferation and death (12). Particularly, the “competing endogenous RNA (ceRNA) hypothesis” has been suggested, which states that many lncRNAs act as ceRNAs by sequestrating target miRNAs; abundant lncRNAs harboring similar microRNA target sequences can sequester cytoplasmic miRNAs, thus regulating miRNAs-target genes indirectly (13–15). lncRNAs have been shown to maintain cellular homeostasis and enable adaptive survival in hypoxia-associated cancer processes (7,16–18). However, the specific roles of lncRNAs in HPH remain poorly understood.

One lncRNA involved in the response to hypoxia is maternally expressed gene 3 (lncRNA-MEG3), which is increased during exposure to hypoxic conditions (19). Elevated expression of lncRNA-MEG3 has also been detected in the developing otocyst and differentiated neural cells (20). Numerous studies have demonstrated that lncRNA-MEG3 is a tumor suppressor (21). lncRNA-MEG3 expression is lost in multiple cancer cell lines of various tissue origins and its over-expression inhibits tumor cell proliferation and cell cycle progression (22–24). These controversial observations support a role for lncRNA-MEG3 in cellular homeostasis and adaptive survival under hypoxic stress conditions depending on the tissue/cell type. Particularly, using a computational prediction approach, we predicted that lncRNA-MEG3 harbors miR-328 target sequences (nucleotides (nt) 2071–2094). Thus, lncRNA-MEG may be involved in the development of HPH by interacting with miR-328.

In the present study, we examined the expression of lncRNA-MEG3 in pulmonary arteries (PAs) in a hypoxia-induced PH animal model and PASMCs. Upon hypoxia, upregulation of lncRNA-MEG3 acts as a ceRNA to suppress miR-328 expression, leading to the expression of insulin-like growth factor 1 receptor (IGF1R), and excessive proliferation of PASMCs. Accordingly, lncRNA-MEG3 is highly expressed in PASMCs of idiopathic PAH (iPAH) patients, suggesting lncRNA-MEG3 as a new target for pharmaceutical and the treatment of PH.

## MATERIAL AND METHODS

### Animal Model of Chronic Hypoxia-Induced PAH and Tissue Preparation

Adult male C57BL/6 mice (mean weight of 30 g) were obtained from the Experimental Animal Center of Harbin Medical University, which is completely accredited by the policies of Institutional Animal Care and Use Committee. The animal study was approved by the ethics review board of Harbin Medical University ([2012]-006). Mice were randomized for exposure for the indicated times to normal and hypoxic environments with fractional inspired oxygen (FiO2) of 0.21 and 0.12, respectively. After the indicated hypoxia exposure period, mice were anesthetized with chloral hydrate (40 mg/kg, i.p.). Right ventricular systolic pressure (RVSP) and right ventricular to left ventricular + spatial weight ratio (RV/LV+S), two indices of pulmonary artery systolic pressure, were measured to confirm the success of producing the HPH model. Then the organs were excised for the following experiments. For the SUGEN models, Animals received Su5416 (20 mg/kg subcutaneously; dissolved in DMSO; Sigma - Aldrich, St Louis, MO, USA), immediately followed by hypoxia exposure immediately. After 3 weeks of hypoxia, animals were returned to room air for another 4 weeks.

**Histological and morphometric analyses, Cell Culture, RNA Isolation and Real-time Polymerase Chain Reaction (Real-time PCR), protein preparation and western blot analysis, Immunofluorescence assay, Cell cycle progression analysis, Cell viability assay, BrdU assay**

These have been described in the Online Data Supplement.

### Fluorescence In Situ Hybridization

To confirm the cellular localization of lncRNA-MEG3 in PASMCs, fluorescencein situ hybridization (FISH) was conducted; lncRNA-MEG3 FISH Probe Mix was designed and synthesized by RuiBo biology Co. Ltd. (#C10910, Guanzhou, China). Experiments were performed according to the manufacturer’s protocol. Briefly, excised organs were embedded in paraffin and 4-μm cryosections were deparaffinized. Following rehydration in phosphate-buffered saline and acetylation, the slides were hybridized at 23°C for 3 h and then incubated with 200 ng custom-designed digoxigenin-labeled lncRNA-MEG3 probes (probe sequences: 5’-GGTCCCTCTCTGGCAACTGTTCATTCATTTGATGC-3’, 5’-CTCAGGCCTGTCGCGTCTTCCTGTGCCATTTGCTG-3’, 5’-TTTACAAATGGACTCTTGTTCCGTCGTTTCCCACC-3’) in 200 μL hybridization buffer) at 60°C overnight. Following sequential washing in 5x saline sodium citrate (SSC), 0.2×SSC (20×SSC contains 3 M NaCl, 0.3 M Na-citrate, pH 7.0), and Tris-buffered saline (TBS), the slides were blocked using 10% fetal calf serum in TBS for 2 h and incubated with alkaline phosphatase-labeled anti-digoxigenin antibodies (3.75 U/mL; Roche, Basel, Switzerland) in TBS at 4°C overnight. After washing with TBS, bound antibodies were detected by microscopy. Probes targeting alpha-smooth muscle actin (αSMA) mRNA (5’-CCTTGAAGTACCCGATCGAACATGGCATCA-3’; 5’-CTGACTACCTCATGAAGATCCTGACTGAGC-3’; 5’-GGAATCTGCTGGCATCCATGAAACCACCTA-3’) was used as positive controls. PBS without probes was used as negative control.

For in vitro cultured cell FISH, PASMCs were fixed with 4% paraformaldehyde, and then incubated with lncRNA-MEG3 FISH Probe Mix for overnight at 37°C, Fluorescence images were obtained with a fluorescence microscope. U6 FISH Probe Mix was used to localize the nuclei, 18S FISH Probe Mix was used to localize the cytoplasm, and DAPI (4,6-diamidino-2-phenylindole) was used to label the nuclei. Images were captured by microscopy and analyzed with Image software (Image Pro Plus).

### lncRNA-MEG3 Knockdown In Vivo and In Vitro

To silence the expression of lncRNA-MEG3, PASMCs were transfected with the small interfering RNA (siRNA) targets to lncRNA-MEG3 (siMEG3-936, 5’-GCGUCUUCCUGUGCC AUUUTT-3’ sense; 5’-AAAUGGCACAGGAAGACGCTT-3’ antisense), (siMEG3-1147, 5’-CC UCCUGGAUUAGGCCAAATT-3’ sense; 5’-UUUGGCCUAAUCCAGGAGGTT-3’ antisense) Non-targeted control siRNA (siNC, 5’-UUCUCCGAACGUGUCACGUTT-3’ sense; 5’-ACGUG ACACGUUCGGAGAATT-3’ antisense), using X-treme Gene siRNA Transfection Reagent (Roche). In some experiments, lncRNA-MEG3 was knocked down with gapmers (gapmer-1, TCCATTTGCCTCATAA; gapmer-3, CACTCCATCACTCATA), which was designed and synthesized by Exiqon (Vedbaek, Denmark). To knock down the expression of lncRNA-MEG3 in pulmonary arteries (PAs), siRNA against lncRNA-MEG3 (siMEG3) was loaded with octaarginine (R8) conjugated PEG2000-Lipid (R8-Lip) to form a lung-specific delivery system (R8-Lip-siMEG3) as previously described with some modifications.(25) Briefly, octaarginine (R8) and DSPE-PEG2000-Mal (molar ratio: 1.5:1) were mixed in chloroform/methanol (v/v = 2:1) at room temperature with gentle stirring for 48 h. The mixture was evaporated under vacuum and then re-dissolved in chloroform, after discarding the insoluble material; the supernatant (DSPE-PEG2000-R8) was evaporated again under vacuum. Lipid compositions of the prepared liposomes were as follows: R8 modified liposomes (R8-LIP), SPC/Chol/DSPE-PEG2000/DSPE-PEG2000-R8 (molar ratio 59:33:3:5). A lipid film was produced by rotary evaporation of all lipids in chloroform. The films were left under vacuum for 2 h. Hydration buffer was added to produce a concentration of 10 μmol/3 mL of lipid. The lipid suspensions were sonicated with a probe sonicator at 80 W for 2 min. siMEG3 and lipid ingredients, at a siRNA to lipid molar ratio of 1:40, were dissolved in chloroform to prepare siRNA-loaded liposomes. Free siRNA was removed using a Sephadex-G50 column and the pellets were collected. The bio-distribution of R8-Lip was evaluated with a labeled siRNA (si-MEG3) and fluorescence images were captured. Additionally, αSMA stains or CD31+ stains were used to detect smooth muscle and endothelial compartments with delivery. To efficiently knock down the expression of lncRNA-MEG3, R8-Lip-siMEG3 was repeatedly injected on days 1, 4, 7, and 14 during experiment. R8-Lip and siMEG3 plus Lipofectamine 2000 (Life Technologies, Carlsbad, CA, USA) were used as controls and repeatedly injected. In some experiments, AMO-328 (miR-328 specific 2-O-methyl antisense inhibitory oligoribonucleotide) was loaded into the liposome together with siMEG3. The efficiency of MEG3 knockdown by R8-Lip-siMEG3 was confirmed by using Real-Time PCR and FISH.

### lncRNA-MEG3 Overexpression in PASMCs

An over-expression vector for lncRNA-MEG3 was constructed. Briefly, using Mus musculus maternally expressed 3 (Meg3), transcript variant 1, (NR_003633.3) as template, PCR was carried out with TGAACCGTCAGATCGAATTCGCCACCGGGGACCTTTACAGACC and AATCCAGAGGTTGAGGATCCTTACAGGTGCACCTAATTGGATGGATA to obtain a fragment of 1644 base pairs (bp), which was the gene sequence of lncRNA-MEG3 and upstream and downstream recombination exchange arms. The pLenti-EF1a-EGFP-P2A-Puro-CMV-MCS (H145) vector (CutSmart, New England Biolabs, Ipswich, MA, USA) was digested with EcoR I-HF and BamH I-HF, and the carrier was recovered by electrophoresis after scanning for the 8.8–kb fragment. Enzyme digestion reactions contained 2 μg plasmid, 1 μL enzyme, 5 μl buffer and double-distilled water to a volume of 50 μl; reactions were incubated at 37°C for 2 h. After 1% agarose gel electrophoresis, the vector was removed from the gel and recovered with a gel recovery kit. The lncRNA-MEG3 PCR product was ligated into the vector using a seamless cloning kit to obtain the target plasmid with ampicillin resistance. PASMCs were transfected with the plasmid, and then harvested for RNA isolation, cell proliferation assay, cell migration assay, and flow cytometry analysis.

### MicroScale Thermophoresis (MST)

Sample Preparation: For the experiment FAM-labeled ssRNA of lncRNA-MEG3 molecules was amplification by (5’-GTTTCTGGGCTACGGGTTTG-3’ sense; 5’-CATACTGTTGTCACTCACCC-3’ antisense). The PCR product of ssRNA is 163 nt (2030–2193). Unlabeled single chain miR-328 mimics (5’-CUGGCCCUCUCUGCCCUUCCGU-3’) or control negative control (5’-UUGUACUACACAAAAGUACUG-3’) was added in 1:1 dilutions. For the measurement the samples were filled into standard capillaries. The measurements were conducted on a NanoTemper Monolith NT.115 instrument. The Analysis was performed at 22 % LED power and 20 % MST power, “Fluo. Before” time was 5 seconds, MST on time was 30 seconds and “Fluo. After” time was 5 seconds.

### Statistical Analysis

Quantitative data are expressed as the means ± SEM. Data analysis was performed with paired Student’s t test (for two means) or one-way analysis of variance followed by Dunnett’s test (for >2 means) where appropriate. No statistical method was used to predetermine sample size. Sample sizes (n) were reported in the corresponding figure legend. Significance levels of P < 0.01 (**) and 0.05 (*) were considered statistically significant.

## RESULTS

### Hypoxia Increases the lncRNA-MEG3 Expression

To determine the expression pattern of lncRNA-MEG3 in the development of HPH, male mice were exposed to hypoxia (10% FiO2) for 21 days. RVSP and RV/LV+S, two indirect indicators of pulmonary artery hypertension (Supplemental Figure S1A and S1B), were significantly higher in the hypoxic group than in normal controls. Moreover, hematoxylin-eosin (H&E) staining (Supplemental Figure S1C) demonstrated that the morphology of pulmonary vascular remodeling, as the thickness of vessel wall to vessel diameter ratio was significantly higher in hypoxia models. Therefore, the murine model of chronic HPH was validated in our study.

Next, we detected lncRNA-MEG3 expression in isolated pulmonary arteries (PAs) from HPH mice by using FISH, which revealed that lncRNA-MEG3 (green color) was significantly up-regulated by hypoxia at 21 days (Figure 1A). As a positive control, probes targeting αSMA mRNA was used (red color); the data also revealed increased expression of αSMA after hypoxia exposure (Figure 1A). We also examined the time-dependent induction of lncRNA-MEG3 expression under hypoxic conditions. FISH results revealed that lncRNA-MEG3 was upregulated by hypoxia with significant induction at 4 days and maximal induction at 8–12 days (Figure 1B). Moreover, real-time PCR confirmed that lncRNA-MEG3 was potently induced by hypoxia in a time-dependent manner in isolated PAs (Figure 1C). We also detected lncRNA-MEG3 expression in other tissue (heart, lung, spleen, liver and kidney) and the isolated aorta and carotid arteries, the results showed increasing lncRNA-MEG3 expression only presented in PAs, but decreased or unchanged expression in other tissues (Supplemental Figure S2). Hypoxia induced lncRNA-MEG3 expression in PASMC prompted us to further examine lncRNA-MEG3 expression in PASMC of idiopathic pulmonary arterial hypertension (iPAH). Real-time PCR consistently showed that the expression of lncRNA-MEG3 was higher in PASMCs of iPAH patients than in normal PASMCs (Figure 1D). To further confirm the effect of hypoxia on lncRNA-MEG3 expression, two cells lines - human pulmonary artery smooth muscle cells (hPASMCs) (Figure 1E, left panel) and mouse pulmonary artery smooth muscle cells (mPASMCs) (Figure 1E, right panel) were exposed to 3% FiO2 to induce hypoxia, real-time PCR confirmed that hypoxia augmented lncRNA-MEG3 expression in a time-dependent manner.

**Figure 1.**
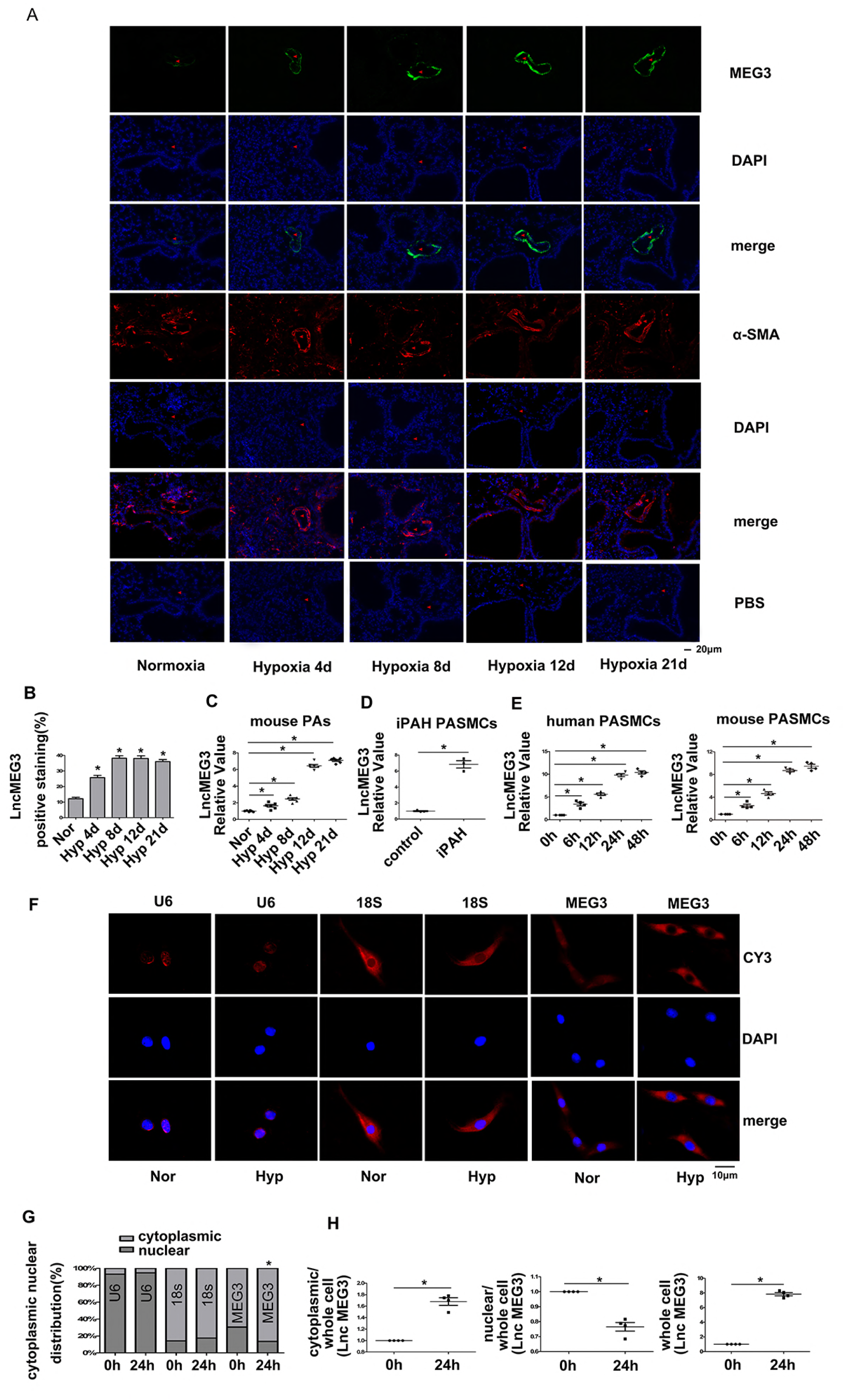
Effects of hypoxia on long noncoding RNA (lncRNA)-maternally expressed gene 3 (MEG3) expression. **A**, Mouse was exposed to hypoxia (10% FiO_2_) for indicated times, lung section was used to detected lncRNA-MEG3 expression by fluorescence in situ hybridization (FISH). FISH probes targeting to αSMA mRNA were used as positive control, PBS was used as negative control. **B**, Bar graph showed that hypoxia time-dependently increased the lncRNA-MEG3 expression in mouse pulmonary arteries (PAs) (n = 3). **C**, real-time PCR confirmed that upregulation of lncRNA-MEG3 by hypoxia (n = 6). **D**, Higher expression of lncRNA-MEG3 was detected in PASMCs of human idiopathic pulmonary artery hypertension patients (n = 3). **E**, Hypoxia increased lncRNA-MEG3 expression in cultured pulmonary artery smooth muscle cells (PASMCs) from human (left panel) and mouse (right panel). **F**, FISH showing the distribution lncRNA-MEG3 signal (red, CY3 staining) in the nucleus (blue, stained with 4,6-diamidino-2-phenylindole) and cytoplasm. U6 and 18S RNA were used as controls for the localization of the nucleus and cytoplasm, respectively. **G**, Quantification of fluorescence showing that hypoxia induced changes in the distribution of lncRNA-MEG3 in the nuclear and cytoplasmic fractions. The graph represents the means ± SEM of cytoplasmic and nuclear fluorescence signals from >50 cells in 3 independent experiments. **H**, Isolation of cytoplasm and nucleus following by real-time PCR data confirmed that hypoxia increased lncRNAs-MEG3 expression, which was mainly located in the cytoplasm but not in the nucleus. Data represent means ± SEM from indicated independent experiments. Student’s t test (for two means) or one-way analysis of variance ANOVA followed by Dunnett’s test (for >2 means). *P < 0.05, **P < 0.01.

To examine the subcellular localization of lncRNA-MEG3, we evaluated lncRNA-MEG3 expression in cytoplasmic versus nuclear fractions using lncRNAs FISH probes labeled with CY3 (red color). As a control, a probe targeting to U6 or 18S RNA was used as a marker of the nucleus or cytoplasm, respectively. LncRNA-MEG3 was detected in both the cytoplasm and nucleus of PASMCs under normoxia conditions. After cells were exposed to hypoxia (3% FiO2) for 24 h, lncRNA-MEG3 was still highly enriched in the cytoplasm, but decreased in the nucleus of PASMCs (Figure 1F). LncRNA-MEG3 distribution in cytoplasm was increased from 69.46 % ± 2.47 % to 86.5 % ± 2.55 %, yet decreased from 30.54 % ± 2.47 % to 13.5% ± 2.55% in nucleus (Figure 1G). However, the localization of U6 and 18S RNA were not significantly changed by hypoxic conditions (Figure 1G). Moreover, by isolating the cytoplasmic and nuclear RNA, followed by real-time PCR (Figure 1H), we confirmed that lncRNA-MEG3 expression was mainly increased in the cytoplasm, but decreased in the nucleus, indicated lncRNA-MEG3 translocated from the nucleus to the cytoplasm during hypoxia exposure.

Because hypoxia-inducible factor (HIF) transcription factors play an important role in the hypoxia response, we examined whether lncRNA-MEG3 expression is regulated by HIF. HIF isoforms were knocked down by siRNA targeting HIF1α, HIF1β, HIF2α and HIF2β, we found that the increased expression of lncRNA-MEG3 by hypoxia or cobalt chloride was markedly blocked by siHIF1α and alleviated by siHIF1β, but not by siHIF2α or siHIF2β (Supplemental Figure S3A). As a positive control, cobalt chloride (CoCl2, known activators of HIF) significantly evoked expression of lncRNA-MEG3 was attenuated siHIF1α and alleviated by siHIF1β (Supplemental Figure S3B). To complement the results from the knockdown of HIF isoforms, we also examined the effects of HIF isoforms overexpression on lncRNA-MEG3 expression. Consistently, we found that lncRNA-MEG3 could be potently induced by HIF1α and moderately upregulated by HIF1β plasmid (Supplemental Figure S3C). We also examined the role of histone acetylation in regulating lncRNA-MEG3; the results showed that hypoxia-induced lncRNA-MEG3 expression was prevented by C646 (histone acetyltransferase inhibitor) (Supplemental Figure S3D). Altogether, our results revealed that lncRNA-MEG3 expression was induced by hypoxia in an HIF1-dependent manner (mainly by HIF1α).

### LncRNA-MEG3 Contributes to HPH and PA Remodeling

To determine whether lncRNA-MEG3 is required for PA remodeling and HPH development, we inhibited pulmonary vessel lncRNA-MEG3 using a lung-specific delivery system (R8 peptide conjugated PEG2000-lipid (R8-Lip) which modified siRNA targets to MEG3 (R8-Lip-siMEG3)) in vivo. R8-Lipid-siMEG3 was injected into the tail vein under hypoxic conditions on days 1, 3, 8 and 15, siRNA targets to MEG3 without any modification (siMEG3) and R8-Lip were used as controls. The distribution of R8-Lip targets to the pulmonary smooth muscle layer was confirmed by double-staining of labeled siMEG3 with αSMA, but not between labeled siMEG3 with CD31 (Figure 2A). The cytotoxicity of R8-Lip-siMEG3 was evaluated by MTT assay (Figure 2B), which revealed that R8-Lip-siMEG3 was not cytotoxic towards PASMCs after 24 h. R8-Lip-siMEG3 significantly and efficiently suppressed lncRNA-MEG3 expression (green color), which was confirmed by tissue FISH (Figure 2C-2D) and by real-time PCR (Figure 2E) in isolated PAs from HPH mice. But siMEG3 and R8-Lip did not affect the lncRNA-MEG3 expression under hypoxic conditions. FISH probes targeting αSMA mRNA was used as positive control, PBS was used as negative control (Figure 2C). Increased RVSP and RV/LV+S (Figure 2F) induced by hypoxia were significantly attenuated by R8-Lip-siMEG3, whereas siMEG3 and R8-Lip did not show significant inhibitory effects. Moreover, R8-Lip-siMEG3 inhibited the elevation of proliferating cell nuclear antigen (PCNA) expression (Figure 2G) according to Ki67 staining (Figure 2H). We also measured the muscularization of small pulmonary vessels by Masson trichrome staining, R8-Lip-siMEG3 but not the controls (siMEG3 or R8-Lip) inhibited the increasing percentage of medial wall thickness to the external diameter evoked by hypoxia (Figure 2I). These findings confirm that increased lncRNA-MEG3 leads to thypoxia-induced PA remodeling and HPH development *in vivo*.

**Figure 2.**
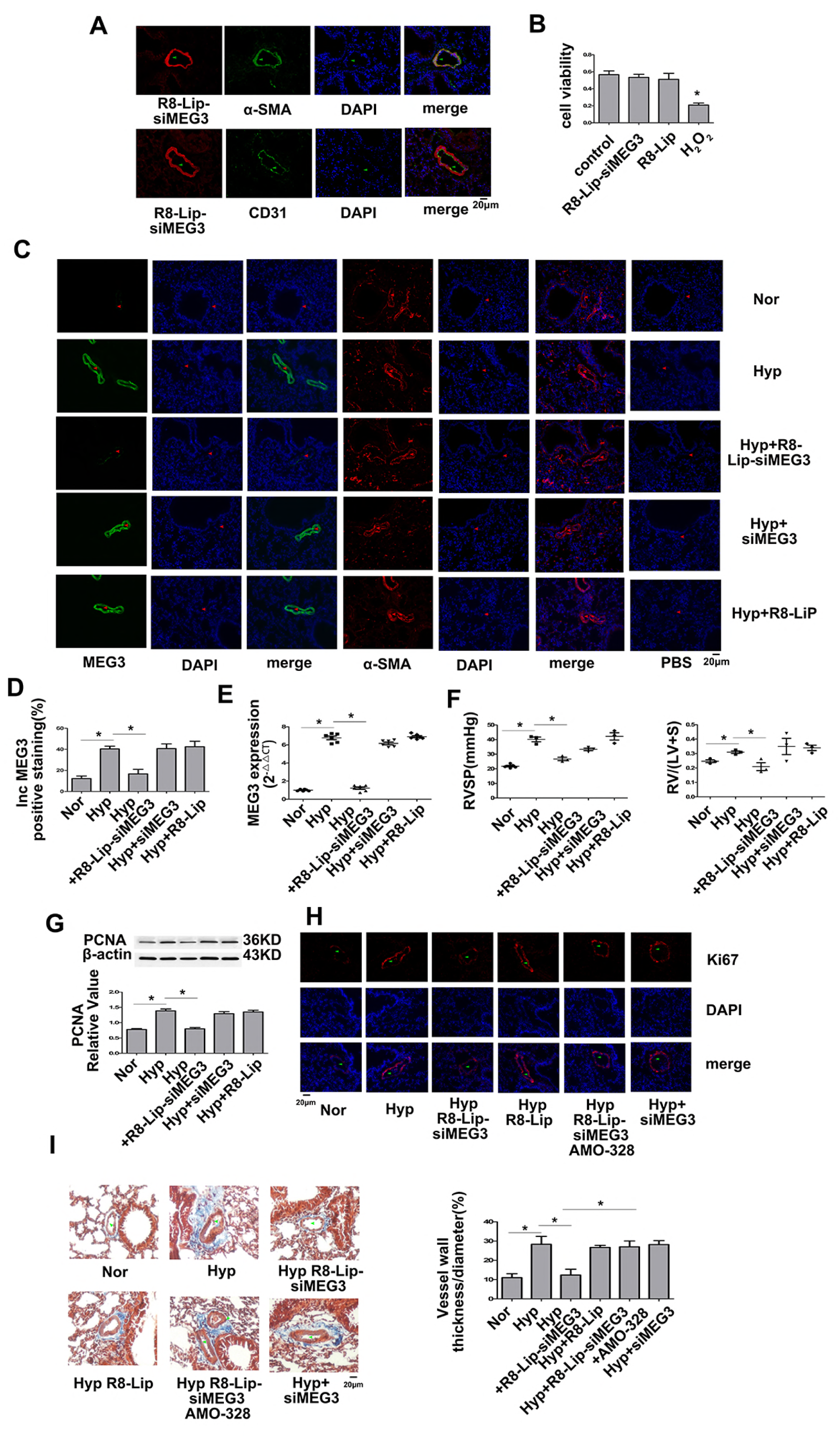
Pulmonary-specific lncRNA-MEG3 knockdown prevented hypoxia-induced pulmonary hypertension. **A**, Colocalization of labeled siMEG3 (red) with immunostaining of αSMA (up panel, green) but not CD31 (lower panel, green) indicated that siRNA was delivered to the smooth muscle layer by the R8-liposome delivery system. **B**, MTT data showed that R8-modified liposomes were not cytotoxic towards PASMCs after 24 h. **C**, Efficiency of R8-liposome-loaded siRNA targets to lncRNA-MEG3 (R8-Lip-siMEG3) on knockdown of lncRNA-MEG3 expression was confirmed by fluorescence in situ hybridization (FISH). FISH probes targeting to αSMA mRNA were used as positive control, PBS was used as negative control. **D**, Bar graph indicated lncRNA-MEG3 positive staining of FISH (n = 3). **E**, real-time PCR in isolated PAs revealed that R8-Lip-siMEG3 significantly reversed the increased lncRNA-MEG3 level induced by hypoxia, whereas non-modified siRNA (siMEG3) and R8-Lip did not show significant effects (n = 6). **F**, RVSP and RV/LV+S, proliferating cell nuclear antigen (PCNA) expression (**G**) and Ki67 staining (**H**) evoked by hypoxia were attenuated by R8-Lip-siMEG3 but not siMEG3 or R8-Lip. **I**, Pulmonary vascular muscularization illustrated by masson trichrome staining. The collagen is blue, the wall thickness percentage of the external diameter in different groups was measured. The results indicated increasing muscularization of small pulmonary vessels by hypoxia, which was attenuated by knockdown of lncRNA-MEG3 by R8-Lip-siMEG3. Data represent means ± SEM from indicated independent experiments. One-way analysis of variance ANOVA followed by Dunnett’s test was used. *P < 0.05, **P < 0.01.

### LncRNA-MEG3 Knock down Inhibits Hypoxia-induced PASMC Proliferation

Because lncRNA-MEG3 was expressed and increased by hypoxia in PASMCs, we further explored the function of lncRNA-MEG3 by knocking down lncRNA-MEG3 expression with siMEG3 or gapmers. Two different siRNAs (siMEG-936 and siMEG3-1417) and gapmers targeting two different sequences were designed to knock down lncRNA-MEG3 expression. All siRNAs and gapmers efficiently inhibited lncRNA-MEG3 expression in a dose-dependent manner (Supplemental Figure S4A). Because no difference was observed in the knockdown of lncRNA-MEG3 expression between the two siRNAs, siMEG-936 was used for further in vivo and in vitro studies. siMEG3 treatment resulted in a reduction of lncRNA-MEG3 levels compared to scramble controls RNAs (negative control, NC). Hypoxia increased the cell viability of PASMCs, and silencing of lncRNA-MEG3 attenuated the effects of hypoxia compared to in NCs (Figure 3A). Indeed, in the analysis of PASMC proliferation, hypoxia markedly enhanced BrdU incorporation (Figure 3B), PCNA expression (Figure 3C), and Ki67 staining (Figure 3D). In the presence of the siMEG3, these effects were reduced nearly to base line levels.

**Figure 3.**
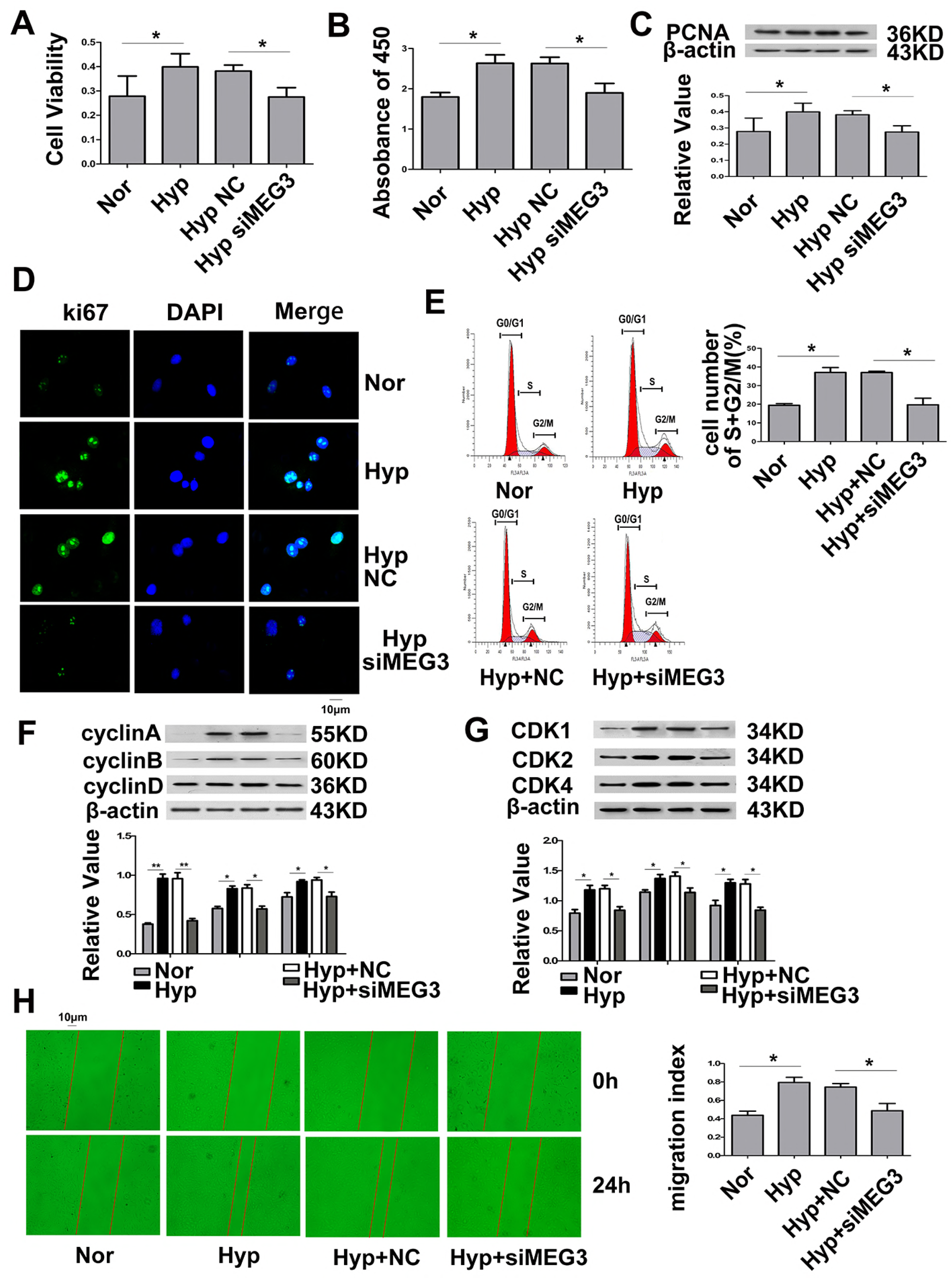
Maternally expressed gene 3 (MEG3) involved in hypoxia-induced PASMC proliferation, cell cycle progression, and cellular migration. PASMCs were transfected with siRNA targeting lncRNA-MEG3 (siMEG3) or negative control nucleotide (NC), and then exposed to hypoxia for 24 h. siMEG3 significantly abolished the increasing PASMC cell viability induced by hypoxia determined by MTT (**A**), BrdU incorporation (**B**), PCNA expression (**C**) determined by immunoblotting, and Ki67 staining (**D**) by immunofluorescence. **E**, Flow cytometry indicated that siMEG3 prevented cell cycle progression. **F**, Cyclin and CDK (**G**) expression were inhibited by siMEG3 under hypoxic conditions. **H**, Cellular migration experiment indicated that siMEG3 reversed hypoxia-induced cell migration. PCNA represents proliferating cell nuclear antigen. CDK represents cyclin-dependent kinases. Data represent means ± SEM from 4 independent experiments. One-way analysis of variance ANOVA followed by Dunnett’s test was used. *P < 0.05, **P < 0.01.

Next, we investigated the function of lncRNA-MEG3 in cell cycle progression. As expected, hypoxia promoted the cell cycle into the S+G2/M phase, whereas lncRNA-MEG3 knockdown attenuated the effects of hypoxia (Figure 3E). Furthermore, the knockdown of lncRNA-MEG3 prevented the increased in the expression of cyclins A, B, and D (Figure 3F), and coordinated the activation of important cell cycle proteins, including cyclin-dependent kinases (CDK1, CDK2, and CDK4) (Figure 3G), which were induced by hypoxia. Moreover, we detected the role of lncRNA-MEG3 in cell migration. The results indicated that lncRNA-MEG3 is involved in increasing cellular migration induced by hypoxia. Knockdown of lncRNA-MEG3 by siRNA inhibited hypoxia-induced migration (Figure 3H). Gapmers of lncRNA-MEGs displayed similar effects on preventing cell proliferation, cell cycle progression, and cellular migration induced by hypoxia (Supplemental Figure S4B–4I).

To confirm the function of lncRNA-MEG3 in hypoxia-induced proliferation and cell cycle progression of PASMCs, we overexpressed one segment of MEG3 (mouse transcript varies 1, nt 1563–3207, 1644 bp) by using a lentiviral vector. As shown in Figure 4, overexpression of MEG3 increased cell proliferation (such as cell viability, BrdU incorporation, PCNA expression, and Ki67 staining), cell cycle progression (flow cytometry, cyclin and CDK expression), and cell migration in the absence of hypoxia. Together, these results suggest that lncRNA-MEG3 is involved in hypoxia-induced PASMC proliferation and cell cycle progression.

**Figure 4.**
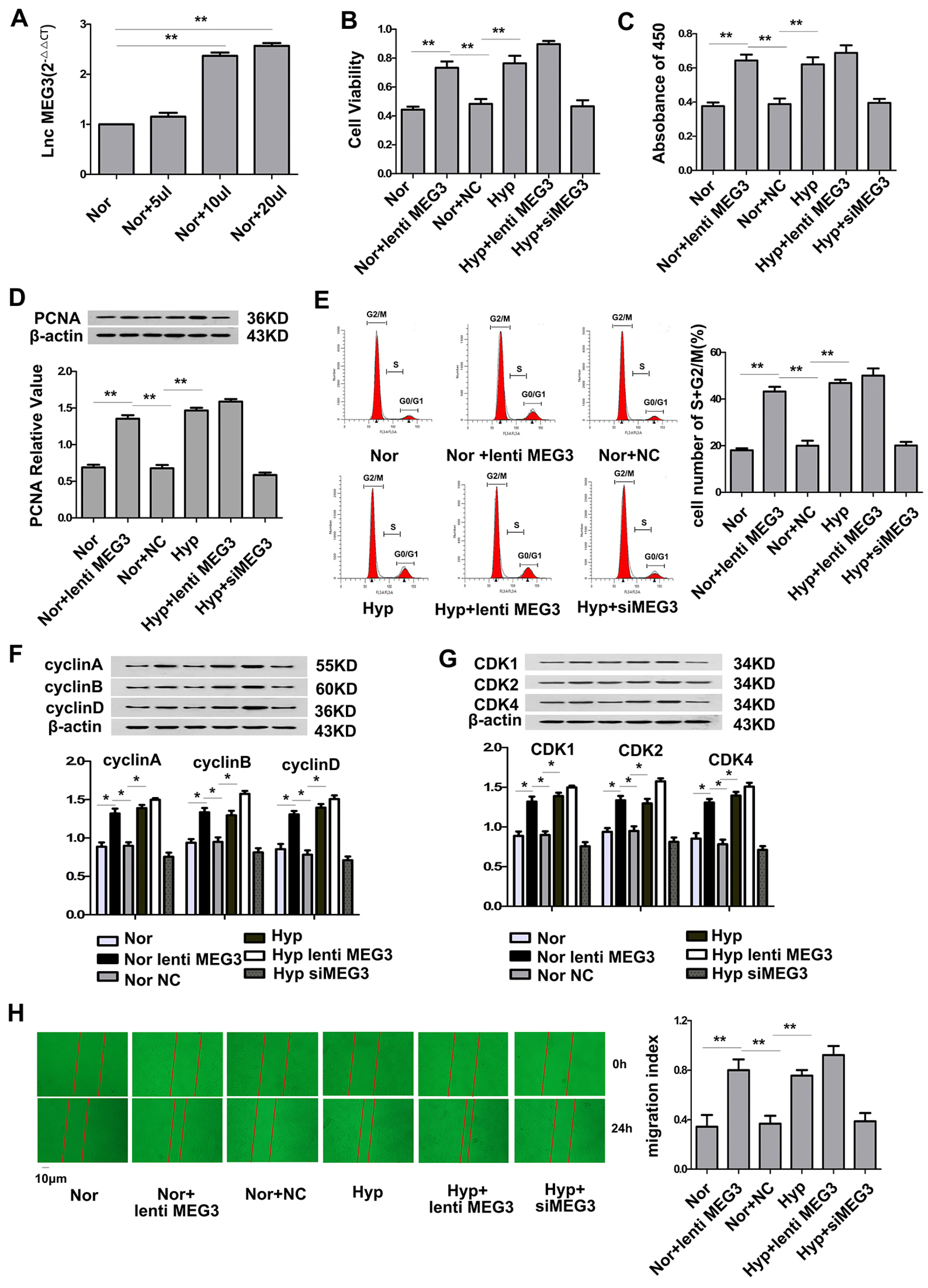
Overexpression of maternally expressed gene 3 (MEG3) induces PASMC proliferation, cell cycle progression, and cellular migration. One segment of lncRNA-MEG3 (mouse transcript varies 1, nt 1563–3207, 1644 bp) was amplified by Real-Time PCR, and then constructed into lentiviral vectors. **A**, Dose-dependent plasmid transfection resulted in enhanced expression of lncRNA-MEG3 in mPASMCs. Overexpression of lncRNA-MEG3 under normal condition resulted in increased cell viability (**B**), BrdU incorporation (**C**), PCNA expression (**D**), cell cycle progression (**E**), cyclin expression (**F**), CDK expression (**G**), and cellular migration (**H**). Data represent means ± SEM from 4 independent experiments. One-way analysis of variance ANOVA followed by Dunnett’s test was used. *P < 0.05, **P < 0.01.

### Association of lncRNA-MEG3 with MicroRNA-328

CeRNA acts as an important mechanism of lncRNA downstream signaling by regulating the expression of miRNA. To understand the molecular mechanism by which lncRNA-MEG3 regulates hypoxia-induced PASMC proliferation and cell cycle progression, we used computational approaches to assess the binding propensity of lncRNA-microRNAs. This analysis identified 5 potential lncRNA-MEG3-interacting microRNAs, including miR-21, miR-30c, miR-138, miR-145, and miR-328 (Figure 5A and Supplemental Figure S5A). To confirme the relationship between these microRNAs and lncRNA-MEG3, we detected the expression of these microRNAs under hypoxic conditions with and without lncRNA-MEG3 knockdown, and found that expression of miR-30c and miR-328 were inhibited, whereas those of miR-138, miR-145, and miR-21 were increased (Figure 5B and Supplemental Figure S5B) by hypoxia. However, among these microRNAs, only miR-328 was regulated by lncRNA-MEG3, as the knockdown of lncRNA-MEG3 reversed the down-regulation of miR-328 under hypoxic conditions (Figure 5B and Supplemental Figure S5C). Furthermore, real-time PCR revealed that miR-328 was decreased in PAs under hypoxic conditions in a time-dependent manner (Figure 5C), and lung-specific knockdown of lncRNA-MEG3 by R8-Lip-siMEG3 rescued this downregulation, but siMEG3 without modification or R8-Lip control did not show effect (Figure 5D). Interestingly, down-regulation of miR-328 rescued by R8-Lip-MEG3 was not detected in lung tissue and right ventricular (Figure 5D). To determine the relationship between lncRNA-MEG3 and miR-328 is applicable in other models of PH, SUGAN/hypoxia mouse models and PASMCs of human idiopathic PAH (iPAH) patients were used. The results revealed that lncRNA-MEG3 was increased, but miR-328 decreased in PAs from SUGEN/hypoxia mouse models (Figure 5E), and in PASMCs of iPAH patients (Figure 5F). Using a computational approach, we found that nt 2071–2094 of lncRNA-MEG3 may directly bindimiR-328. To confirm this hypothesis, we performed a luciferase reporter assay, as shown in Figure 5G. We cloned the putative miR-328 target binding sequence into a luciferase construct; the results showed that miR-328 repressed and AMO-328 (inhibitor nucleotide of miR-328) increased luciferase activity (Figure 5G, left panel). Mutation of the putative miR-328 binding site (marked in red color in Figure 5A) decreased the response to miR-328 (Figure 5G, right panel). In addition, we used MicroScale Thermophoresis (MST) to analyze the interaction of lncRNA-MEG3 with miR-328. The results indicated that miR-328 mimics binding with lncRNA-MEG3 (nucleotide nt 2030–2193) with Kd of 7.46 ± 0.166 μM. As a control, a non-binding mutant miR-328 was used (Kd of 3940 ± 46 μM) (Figure 5H). These results indicate that lncRNA-MEG3 binds to miR-328 in a sequence-specific manner, similar to that by which lncRNAs function as molecular sponges of microRNAs, and up-regulation of lncRNA-MEG3 induced by hypoxia decreased the expression of miR-328 in vivo and in vitro.

**Figure 6.**
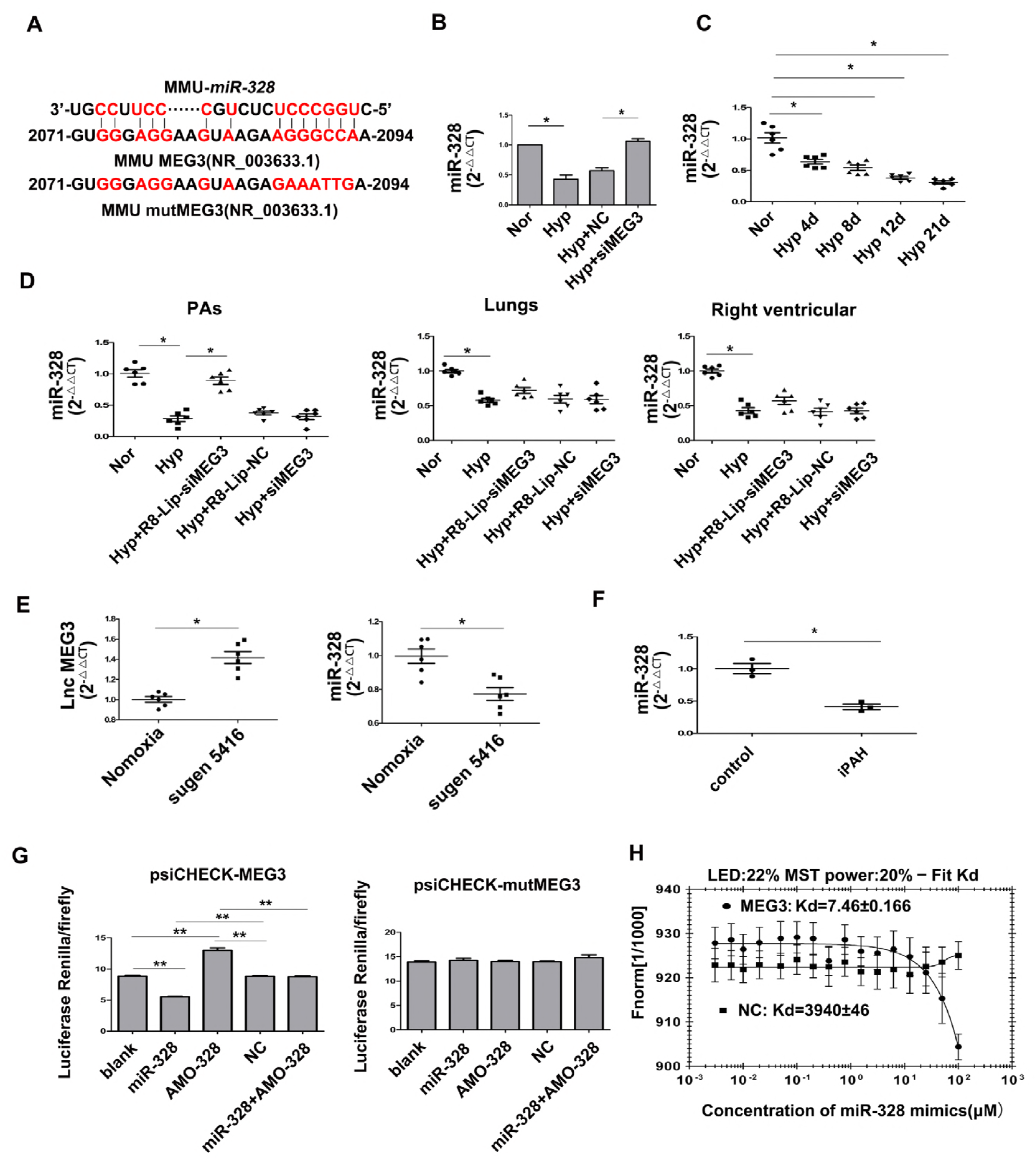
LncRNA-MEG3 acts as ceRNA by sequestrating miR-328. **A**, MiR-328 binding sequences in lncRNAs-MEG3. Red color letters in lncRNAs-MEG3 sequence (lower panel) show mutation sequence used for luciferase experiments. **B**, mPASMCs in culture dish were transiently transfected with 30 nM negative control siRNA (NC) or siRNA against MEG3 (siMEG3), following by hypoxia for 24 h. Downregulated miR-328 by hypoxia was reversed by siMEG3. **C**, Real-time PCR revealed that downregulation of miR-328 during hypoxic conditions with a time-dependent manner (n = 6). **D**, Detection of miR-328 expression in PAs, lung, and right ventricular from hypoxia PAH mouse, and lung-specific knockdown of lncRNA-MEG3 by R8-Lip-siMEG3, siMEG3, and R8-Lip treatment animals, the results indicating that sequestrating of miR-328 by lncRNA-MEG3 occurs in vivo (n = 6). **E**, Detection of lncRNA-MEG3 and miR-328 expression in PAs from SUGAN/hypoxia mouse models. **F**, Lower expression of miR-328 were detected in PASMCs of iPAH patients (n = 3). **G**, mPASMCs were transiently transfected with the indicated agents, followed by transient transfection with 25 ng of psiCHECK plasmid harboring the lncRNA-MEG3 binding region (nt 2071–2094) (left panel) or mutation sequence (right panel). Luciferase activity was assayed 36 h after transfection and normalized to the activity of Renilla luciferase. **H**, MicroScale Thermophoresis (MST) analysis indicated that miR-328 mimics interaction with lncRNA-MEG3 (2030–2193, 163-nt PCR product) with Kd of 7.46 ± 0.166 μM. As a control a negative control nucleotide was used (Kd of 3940 ± 46 μM). Data represent means ± SEM from indicated independent experiments. Student’s t test (for two means) or one-way analysis of variance ANOVA followed by Dunnett’s test (for >2 means). *P < 0.05, **P < 0.01.

### LncRNA-MEG3 functions with its association with miR-328

To investigate the interplay of lncRNA-MEG3 and miR-328 on hypoxia-induced cell proliferation and cell cycle progression, we used miR-328 mimics or inhibitor (AMO-328) to examine the role of lncRNA-MEG3 knockdown on hypoxia-induced events. Both increasing expression of endogenous miR-328 by knockdown of lncRNA-MEG3, and application of exogenous miR-328 mimics reversed the increased cell viability (Figure 6A), BrdU incorporation (Figure 6B), PCNA expression (Figure 6C), and Ki67 staining (Figure 6D), as well as accelerated the cell cycle into the S+G2/M phase (Figure 6E) and increased the expression of cyclins A, B, and D (Figure 6F) and CDK1, CDK2, and CDK4 (Figure 6G) induced by hypoxia. Application of AMO-328 (which disrupts the binding of miR-328 with its targets) reversed the effects of lncRNA-MEG3 knockdown and miR-328 mimics, whereas application of miR-328 mimics during lncRNA-MEG3 knockdown had no additional effects. Moreover, we examined whether anti-miR-328 therapy in vivo by injecting AMO-328 could reverse the effects of lncRNA-MEG3 knockdown. The data showed that anti-miR-328 therapy prevented the increases in expression of Ki67 (Figure 2G) and muscularization of PAs (Figure 2H). We next evaluated the direct targets of the interplay of lncRNA-MEG3 and miR-328. Our previous study showed that miR-328 is involved in HPH by regulating the expression of IGF1R (6). Computational approaches indicated the binding propensity of miR-328 with IGF1R mRNA (Figure 6H). We also observed that IGF1R is upregulated in PASMCs of iPAH patients (Figure 6I). Next, we detected IGF1R expression in mPASMCs, as shown in Figure 6J, the knockdown of lncRNA-MEG3 by siMEG3 or application of miR-328 mimics, or siMEG3 plus miR-328 mimics, all of which inhibited the increase in the expression of IGF1R mRNA and protein levels under hypoxic conditions. Disruption of the binding of miR-328 with AMO-328 reversed the inhibition effects of siMEG3. Moreover, we confirmed IGF1R expression in isolated PAs from HPH mouse models. Up-regulation of IGF1R mRNA and protein levels were prevented by R8-Lip-siMEG3, which was reversed by AMO-328 (Figure 6K). Together, these results demonstrate that lncRNA-MEG3 functions as a ceRNA to sequester miR-328 and increase the downstream target gene IGF1R to regulate cell proliferation, cell cycle progression, and cellular migration during HPH development.

**Figure 6.**
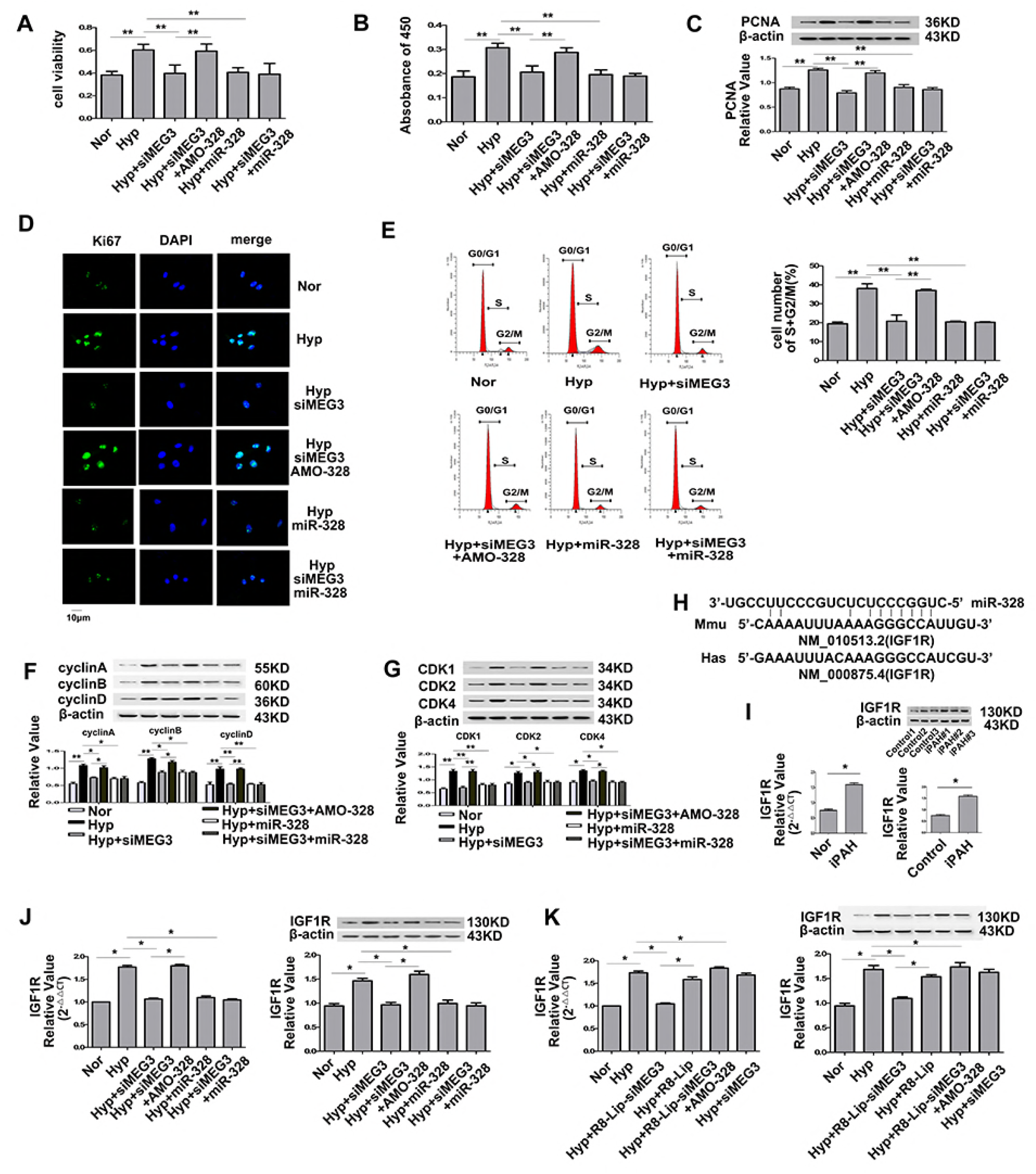
LncRNA-MEG3/miR-328 regulates insulin-like growth factor 1 receptor (IGF1R) expression. Mouse PASMC was transiently transfected with indicated agents, followed by hypoxia for 24 h. Silencing of lncRNA-MEG3 (which results in increased expression of endogenous miR-328), and application exogenous of miR-328 mimics abolished the effects of hypoxia on cell viability (**A**), BrdU incorporation (**B**), PCNA expression (**C**), Ki67 staining (**D**), cell cycle progression (**E**), cyclins A, B, and D expression (**F**), CDK1, CDK2, and CDK4 expression (**G**). IGF1R expression in cultured mPASMCs (**H**) and in isolated PAs (**I**). AMO-328 (miR-328-specific 2-O-methyl antisense inhibitory oligoribonucleotide) reversed the effects of lncRNA-MEG3 knockdown in vitro and in vivo. H, Binding propensity between miR-328 and IGF1R mRNA. **I**, Real-time PCR and western blotting revealed higher expression of IGF1R in PASMCs of iPAH patients (n = 3). Data represent means ± SEM from indicated independent experiments. Student’s t test (for two means) or one-way analysis of variance ANOVA followed by Dunnett’s test (for >2 means). *P < 0.05, **P < 0.01.

## DISCUSSION

In present study, we identified lncRNA-MEG3 as a key regulator of HPH by interacting with miR-328. We showed that lncRNA-MEG3 was up-regulated by hypoxia and involved in hypoxia-induced PASMC proliferation and cell cycle progression. Mechanistically, lncRNA-MEG3 directly binds to miR-328 using nt 2071–2094 as a molecular sponge. Up-regulation of lncRNA-MEG3 sequesters miR-328, leading to increased expression of IGF1R and regulating the development of HPH. This finding implicates lncRNA in HPH and suggests that modulation of the activity of lncRNA-MEG3 and its downstream targets such as miR-328 is a novel therapeutic approach for treating this fatal disease.

The most important finding of our study was that lncRNA-MEG3, up-regulated by hypoxia, is involved in hypoxia-induced PASMC proliferation. Although most previous studies showed that overexpression of lncRNA-MEG3 suppresses cell proliferation in multiple cancer cell lines (22–24), our data showed that upregulation of lncRNA-MEG3 is involved in hypoxia PASMC proliferation. Knockdown of lncRNA-MEG3 expression reversed the hypoxia-increased RVSP and RV/LV+S weight ratio. We also showed that silencing of lncRNA-MEG3 attenuated hypoxia-induced PASMC proliferation in PAs which was confirmed by PCNA expression and Ki67 staining. Through knockdown lncRNA-MEG3 in cultured PASMCs, we further showed that hypoxia-induced PASMC proliferation and cell cycle progression occurred in an lncRNA-MEG3 dependent manner. Finally, lncRNA-MEG3 increased in PASMCs of iPAH patients and in SUGEN/hypoxia models. Altogether, these results indicate lncRNA-MEG3 plays an important role in the development of pulmonary hypertension.

Previous studies indicated that down-regulation lncRNA-MEG3 is correlates with hypoxic microenvironment–related cancer metastasis or recurrence (21–24), which depends on the tissue/cell type (26). However, using HPH mice model and in vitro cultured cells, we observed that lncRNA-MEG3 expression was specifically increased by hypoxia in PASMCs, while downregulation of lncRNA-MEG3 expression in all other tissues was detected, even in the lung tissue (Supplemental Figure S2). Our results differed from reports of other diseases, suggesting that the expression of lncRNA-MEG3 induced by hypoxia is specifically elevated in PASMCs and that upregulation of lncRNA-MEG3 contributes to PASMC proliferation and cell cycle progression. In addition to tissue-specific distribution, our study also showed that lncRNA-MEG3 was primarily localized to the cytoplasm of PASMCs. These results are consistent with those of a previous study showing that lncRNA-MEG3 retains its cytoplasmic subcellular localization in otocysts undergoing proliferation (20). The results suggested that the effect of lncRNA-MEG3 on PASMC proliferation is not related to the cellular location of lncRNA-MEG3 in pulmonary hypertension, but is dependent on the expression of lncRNA-MEG3. Based on our experimental results, lncRNA-MEG3 was only increased by hypoxia in PASMCs, and that alternative mechanism of the lncRNA-MEG3 regulation of PASMC proliferation greatly differs from those in other diseases, even in hypoxic microenvironment-related diseases. The specific effect of lncRNA-MEG3 on PASMCs may depend on the lung vascular responses to hypoxia. Further studies are needed to evaluate these alternative effects among PASMCs and other tumor cell lines.

We observed an increase in lncRNA-MEG3 in hypoxia PAs and PASMCs. The precise factors regulating lncRNA-MEG3 expression by hypoxia remain unclear. A previous study indicated that the regulation of lncRNA-MEG3 by miR-29a was methylation-dependent (24). Cyclic AMP stimulates MEG3 gene expression in cells through a cAMP-response element site in the promoter region (27), suggesting that an alternative regulation pathway functions in PASMCs during hypoxia. By analyzing the promoter of lncRNA-MEG3, we found there are multiple binding sites for transcript factors, such as NFAT2, STAT3, and HIF. Our data indicated that lncRNA-MEG3 was induced by hypoxia in an HIF1-dependent manner (primarily HIF1α) (Supplemental Figure S4). The role of HIF in oxygen-dependent regulation of lncRNA-MEG3 in vivo remains to be elucidated. LncRNA-MEG3 expression was only induced in the PAs but not in other organs, which may be a lung system-specific response to hypoxia; for example, the pulmonary artery contracts but systemic vasculature dilates in response to hypoxia. Indeed, whether other transcript factors and epigenetic regulatory factors, such as DNA methylation or histone acetylation, are involved in manipulating the lncRNA-MEG3 expression during hypoxic conditions in PASMCs requires further analysis. Moreover, lncRNA-MEG3 is a sexually dimorphic gene, showing significant female bias with expression 1.36-fold higher expression in the female cortex/hippocampus than in males (28). Whether the different response of lncRNA-MEG3 to hypoxia varies between females and males, and such sex-dependent expression in HPH cannot be excluded and therefore will be pursued in future studies.

The interaction between lncRNAs and microRNAs has been shown to play important roles in diverse biological processes. Our previous study demonstrated that miR-328 is downregulated in the PAs of experimental animals under hypoxic conditions (6). The present study provided insight into the regulation of miR-328 expression. Notably, we showed that lncRNA-MEG3 uses nt 2071–2094 to bind miR-328, leading to the downregulation of miR-328. This was confirmed by following findings: first, hypoxia inhibited miR-328 expression was attenuated by lncRNA-MEG3 knockdown; second, the luciferase reporter assay indicated that lncRNA-MEG3 bound to miR-328 in a sequence-specific manner; third, MicroScale Thermophoresis results confirmed the interaction of lncRNA-MEG3 with miR-328; fourth, knockdown of lncRNA-MEG3 inhibited the expression of IGF1R (miR-328-regulated gene), and finally, increased lncRNA-MEG3, decreased miR-328, and increased IGF1R were presented in PASMCs of iPAH patients. Although we excluded four other microRNAs displaying highly binding propensity with lncRNA-MEG3, it would be interesting to determine whether lncRNA-MEG3 binds to other microRNAs or proteins. Specifically, lncRNA-MEG3 is known to affect the P53 pathway and transforming growth factor β pathways (29), so it is important to determine whether this association is involved in PASMC proliferation under hypoxic conditions.

The delivery system of R8-modified liposomes has shown potential as a pulmonary drug delivery system for PAH treatment (25). We modified the lung-specific delivery system using siRNA-loaded liposomes (R8-Lip-siMEG3) and found that the biodistribution of R8-Lip in the lung reached high levels in 1 h, which were maintained for 24 h. Thus, we repeated this application of R8-Lip-siMEG3 during hypoxia treatment. Real-time PCR and FISH confirmed the knockdown efficiency of this approach, while non-modified siRNA and R8-Lip showed no significant effects (Figure 2A and 2B). Moreover, RVSP and RV/LV+S indirect indicated the efficiency of R8-Lip-siMEG3 in preventing an increase in pulmonary artery pressure (Figure 2C and 2D). Therefore, using R8-Lip-siMEG3 for knockdown of lung lncRNA-MEG3 expression may provide a foundation for designing new mechanism-based therapies for HPH.

The present study revealed the involvement of lncRNA-MEG3 in HPH, which functions as a molecular sponge for miR-328. Through additional studies of how lncRNAs functions in various diseases (30), lncRNAs may become a new pharmaceutical target and treatment target of pulmonary hypertension.

## AVAILABILITY

None

## ACCESSION NUMBERS

None

## SUPPLEMENTARY DATA

Supplementary Data are available at NAR online.

## ACKNOWLEDGEMENT

We would like to thank Guangtian Wang for the preparation of R8-Lip-siMEG3. The PASMCs of iPAH patients were provided by Professor Jian Wang from State Key Laboratory of Respiratory Diseases, Guangzhou Institute of Respiratory Disease, and The First Affiliated Hospital of Guangzhou Medical University.

## FUNDING

This work was supported by the National Natural Science Foundation of China [31771276 and 31471095 to D. Zhu; 31701010 to Y. Xing; and 31500936 to X. Zheng]; and Specialized Research Fund for the Doctoral Program of Higher Education [20112307110022 to D. Zhu]; and the Natural Science Foundation of Heilongjiang Province [C2016038 to Y. Xing; and C2017042 to X. Zheng]; and China Postdoctoral Science Foundation [2016M601452 to Y. Xing]; and Postdoctoral Foundation of Heilongjiang Province [LBH-Z16239 to Y. Xing]; and Key Laboratory of Cardiovascular Medicine Research (Harbin Medical University), Ministry of Education [2013008 to Y. Xing]; and Postdoctoral career development Fund of Heilongjiang Province [LBH-Q14119 to X. Zheng]; and Harbin medical University Scientific Research Innovation Fund [2016JCZX05 to Yan Xing, and 2017JCZX09 to X. Zheng]; and Foundation of Daqing Department of Science and Technology [Szdy-2015-02 to Y. Xing].

Funding for open access charge: National Natural Science Foundation of China.

## CONFLICT OF INTEREST

None

